# A cortex-specific PBP contributes to cephalosporin resistance in *Clostridium difficile*

**DOI:** 10.1101/715458

**Authors:** Yasir Adil Jabbar Alabdali, Peter Oatley, Joseph A. Kirk, Robert P. Fagan

## Abstract

Sporulation is a complex cell differentiation programme shared by many members of the Firmicutes, the end result of which is a highly resistant, metabolically inert spore that can survive harsh environmental insults. *Clostridium difficile* spores are essential for transmission of disease and are also required for recurrent infection. However, the molecular basis of sporulation is poorly understood, despite parallels with the well-studied *Bacillus subtilis* system. The spore envelope consists of multiple protective layers, one which is a specialised layer of peptidoglycan, called the cortex, that is essential for the resistant properties of the spore. We have identified and characterised a penicillin binding protein (PBP) that is required for cortex synthesis in *C. difficile*. Surprisingly this PBP was also found to contribute to cephalosporin resistance, indicating an additional role in the synthesis of vegetative cell wall. This is the first description of a cortex-specific PBP in *C. difficile* and begins the process of unravelling cortex biogenesis in this important pathogen.

## Introduction

*C. difficile* is the most common cause of nosocomial antibiotic-associated diarrhea, with an estimated 453,000 infections and 29,300 deaths per year in the USA alone (Lessa et al., 2015). *C. difficile* infection (CDI) requires prior disruption to the gut microbiota, most commonly due to an administered antibiotic (Smits et al., 2016). As current treatments largely rely on antibiotic therapy, with further consequent damage to the microbiota, recurrent disease is common and is associated with worse patient prognosis (Rupnik et al., 2009). In recent years there have been dramatic changes in *C. difficile* epidemiology, in particular due to the emergence of the epidemic ribotype 027 lineage, a previously rare ribotype that was responsible for a series of large hospital outbreaks in North America in the early years of this century before spreading worldwide (He et al., 2013).

The spore is an absolute requirement for transmission of disease (Deakin et al., 2012), it allows the organism to transit the lethal aerobic environment while also providing significant resistance to desiccation, heat and common disinfectants (Dyer et al., 2019). As a result, the spores shed by an infected individual can survive in the environment for an extended period of time, a particular problem in hospital environments where large numbers of susceptible individuals are housed in close proximity. The process of sporulation is still relatively poorly understood, despite significant advances in recent years (Zhu et al., 2018). We have previously used high-density transposon mutagenesis and TraDIS to identify a subset of *C. difficile* genes required for formation of mature heat-resistance spores (Dembek et al., 2015). In total, transposon insertions in 798 genes were found to significantly impact sporulation, many with no clear homology to previously characterised proteins. Very few of these 798 genes have been studied in *C. difficile* but many have homologues in the well-studied *Bacillus subtilis* sporulation pathway. However, despite the clear parallels between sporulation in *B. subtilis* and *C. difficile*, there are enough critical differences to greatly reduce the value of assumptions based on homology (Paredes et al., 2005; Underwood et al., 2009; Fimlaid et al., 2013). The response regulator Spo0A is the master regulator of sporulation, phosphorylation of which sets in motion a complex asymmetric cell differentiation programme involving the sequential activation of a series of dedicated sigma factors that are in turn responsible for the expression of the individual regulons required for correct spore morphogenesis (Paredes-Sabja et al., 2014). The result is the complex multi-layered spore structure that lends robustness to environmental insult. The spore consists of a dehydrated core surrounded by a membrane and peptidoglycan cell wall (primordial wall) derived from the mother cell envelope. Around this is a thick peptidoglycan cortex, synthesised during spore maturation, and a second membrane, formed as a result of engulfment of the prespore by the mother cell. The outer surface consists of multiple layers of highly crosslinked proteins. The order and timing of synthesis of each of these layers is critical and disruption to any of the steps typically results in the formation of defective spores that lack full resistance properties (Fimlaid et al., 2013).

In *B. subtilis* the peptidoglycan of the primordial wall and cortex differ in structure allowing differentiation by the cortex lytic hydrolases during germination (Gilmore et al., 2004). The primordial cell wall consists of typical alternating β-1→4-linked N-acetyl glucosamine and N-acetyl muramic acid residues, crosslinked by 4-3 linked peptide stems attached to the muramic acid moieties, while in the cortex peptidoglycan every second N-acetyl muramic acid is modified to muramic-δ-lactam, resulting in fewer stem peptides, fewer crosslinks and a more flexible overall structure (Meador-Parton and Popham, 2000). The class B penicillin binding protein (PBP) SpoVD is critical for synthesis of *B. subtilis* cortex (Daniel et al., 1994). During sporulation SpoVD is expressed in the mother cell where it interacts with the SEDS protein SpoVE to enable localisation to the asymmetric division septum (Fay et al., 2010). An N-terminal transmembrane alpha helix anchors the protein in the membrane, placing the majority of the protein in the inter-membrane space where the cortex is ultimately synthesised (Sidarta et al., 2018). The protein consists of a PBP dimerization domain, followed by a transpeptidase domain and a penicillin-binding protein and serine/threonine kinase associated (PASTA) domain, the last of which is dispensable for cortex formation (Bukowska-Faniband and Hederstedt, 2015).

*C. difficile* vegetative cell peptidoglycan is superficially similar to that of *B. subtilis*, albeit with a preponderance of 3-3 cross-linking as a result of L,D-transpeptidase activity (Peltier et al., 2011). The structure of the *C. difficile* cortex peptidoglycan has not been determined, although it is assumed to contain muramic-δ-lactam, and the enzymes required for cortex synthesis have not yet been characterised. Here we set out to identify and characterise the major *C. difficile* cortex PBP.

## Methods

### Bacterial strains and growth conditions

All bacterial strains, plasmids and oligonucleotides used in this study are described in Table 1. *E. coli* strains were routinely grown in LB broth and on LB agar, while *C. difficile* strains were grown in TY broth (Dupuy and Sonenshein, 1998) and on brain heart infusion agar. Cultures were supplemented with chloramphenicol (15 μg/ml), thiamphenicol (15 μg/ml) or cycloserine (250 μg/ml) as appropriate.

**Table 1:**
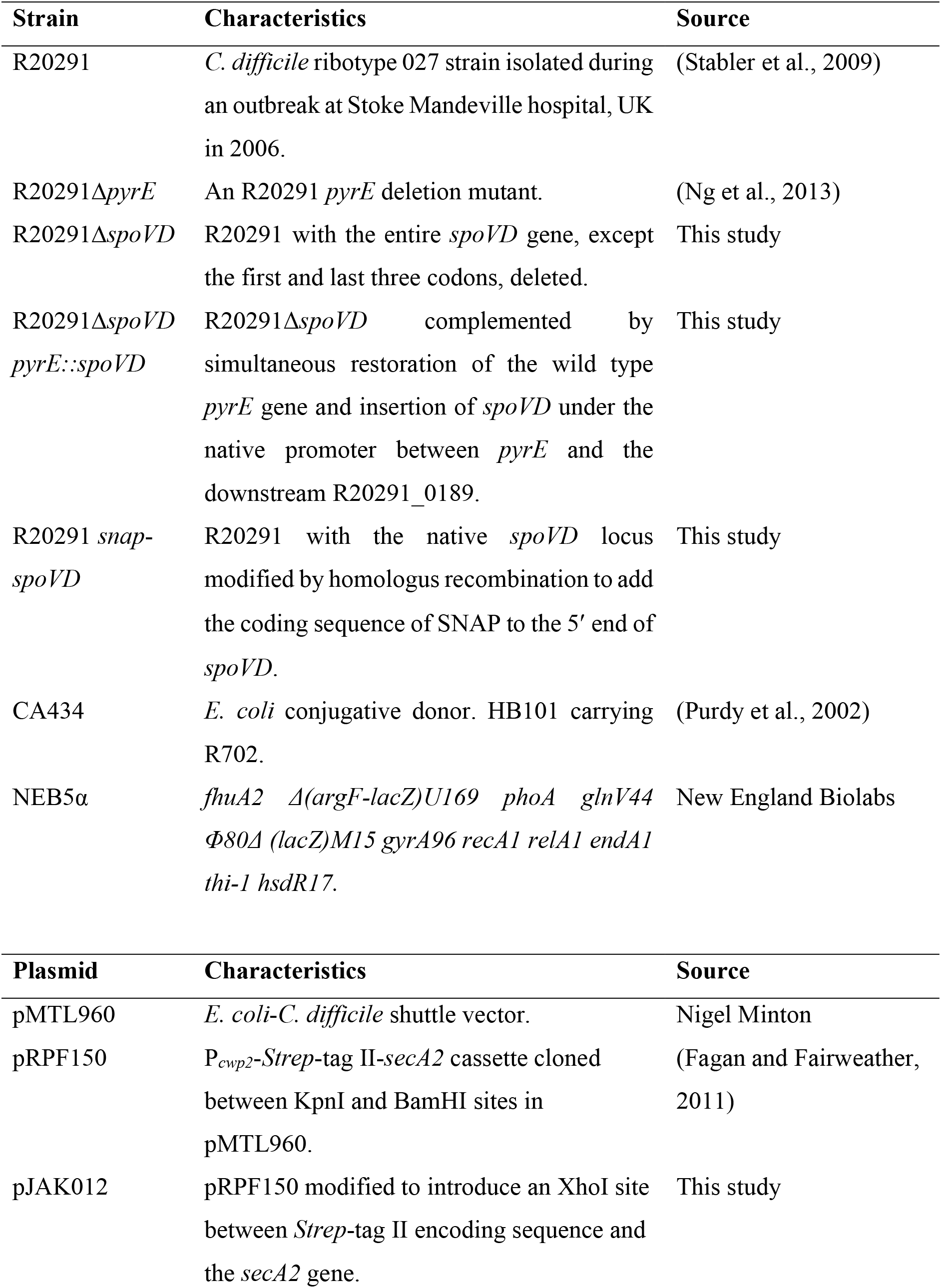
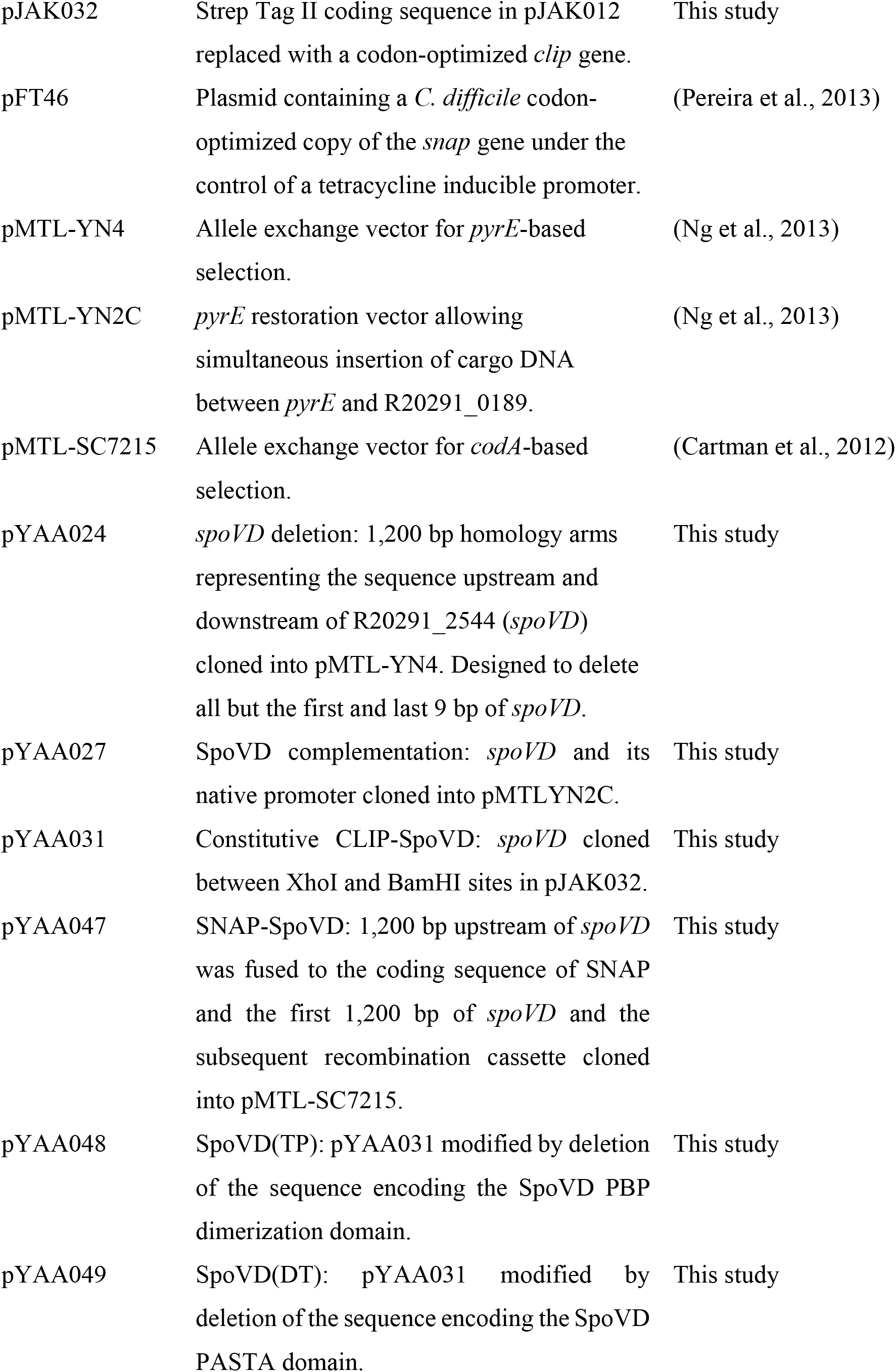
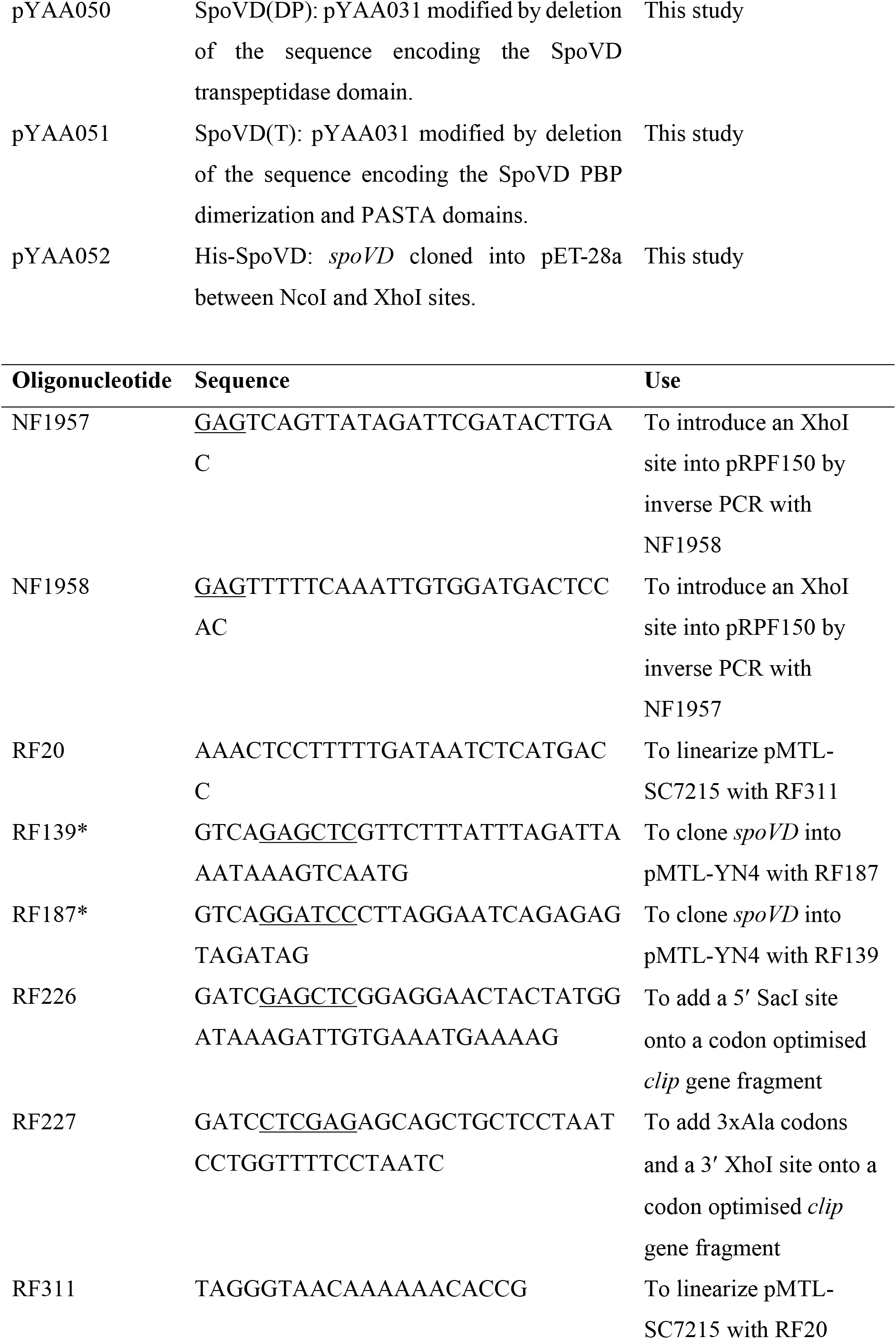
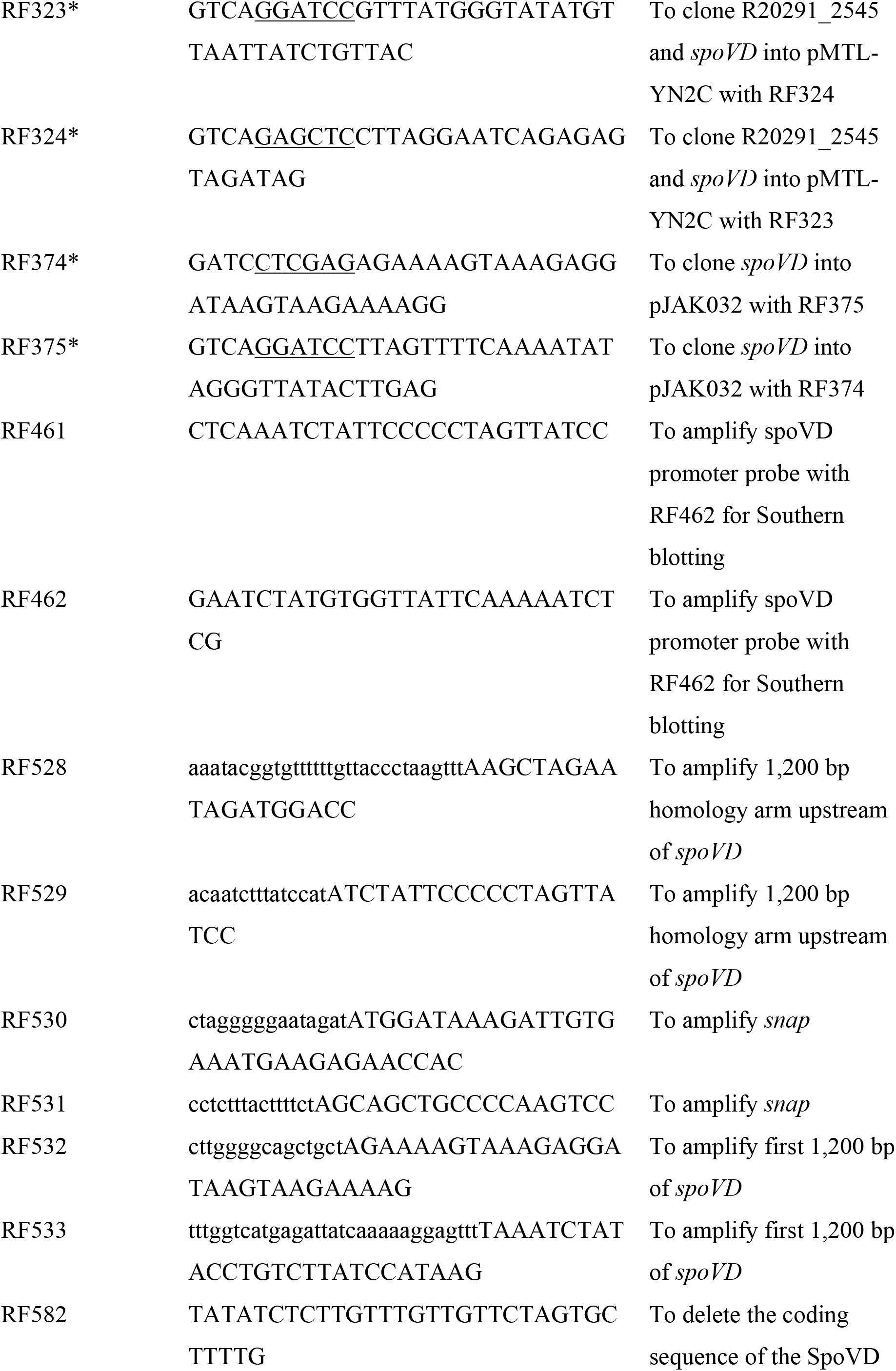
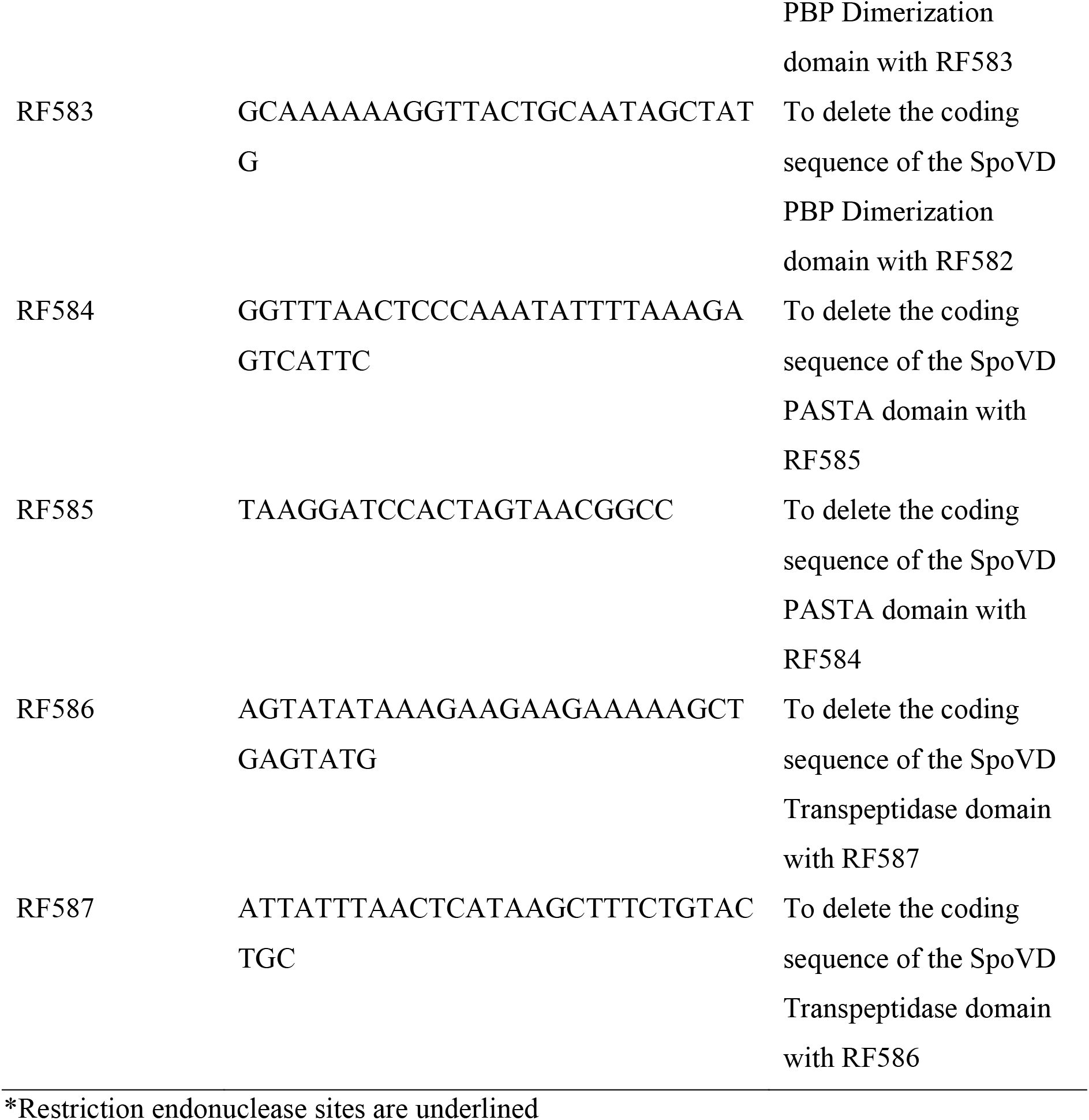
Strains, plasmids and oligonucleotides used in this study

### Molecular biology methods

Routine molecular biology techniques were performed according to the manufacturers protocols except where otherwise stated. PCR using Phusion High-Fidelity DNA Polymerase, plasmid isolation and purification of DNA fragments were performed using kits and reagents supplied by Thermo Fisher Scientific according to the manufacturer’s instructions. Restriction digestion, ligation and Gibson assembly were performed with enzymes supplied by New England Biolabs. Competent *E. coli* were transformed using standard methods and plasmid DNA was transferred to *C. difficile* as described previously (Kirk and Fagan, 2016). *C. difficile* mutants were constructed by homologous recombination as described previously (Cartman et al., 2012; Ng et al., 2013). Mutants were confirmed by PCR and Southern blotting using the Amersham ECL Direct Labelling and Detection System kit (GE) according to the manufacturer’s instructions. A 230 bp probe to the region immediately upstream of *spoVD* was generated by PCD using primer pair RF461/RF462.

### Plasmid construction

pJAK032: pRPF150 was modified by inverse PCR using primer pair NF1957/NF1958 to introduce an XhoI site between the Strep Tag II and SecA2 coding sequences, yielding pJAK012. The Strep Tag II coding sequence was then excised using SacI and XhoI and replaced with a synthetic DNA fragment (IDT gBlock) consisting of a codon-optimized *clip* gene, modified by PCR with primer pair RF226/RF227 to add appropriate SacI and XhoI sites. pYAA024: Homology arms upstream and downstream of *spoVD* were amplified by PCR using oligonucleotide pairs RF68/RF139 and RF69/RF187. The resulting PCR products were joined together in a SOEing PCR reaction and cloned between the BamHI and SacI sites in pMTL-YN4.

pYAA027: *spoVD* expression appears to be driven from a promoter upstream of R20291_2545. In order to ensure complementation at wild type levels a fragment comprising 282 bp upstream of R20291_2545, R20291_2545 itself and *spoVD* was amplified by PCR using primer pair RF324/RF325 and cloned between BamHI and SacI sites in pMTL-YN2C.

pYAA031: *secA2* in pJAK032 was replaced by *spoVD*. *spoVD* was amplified by PCR using primer pair RF374/RF375, digested with BamHI and XhoI and ligated to pJAK032 backbone cut with the same enzymes.

pYAA047: 1,200 bp upstream of *spoVD,* the *snap* tag gene from pFT46 and the first 1,200 bp of *spoVD* were amplified by PCR using primer pairs, RF528/RF529, RF530/RF531 and RF532/RF533 respectively. pMTL-SC7215 was linearized by PCR using primer pair RF20/RF311. The four DNA fragments were then joined in a Gibson assembly reaction.

pYAA048-050: The coding sequence of the SpoVD PBP dimerization domain (pYAA048; primers RF582/RF583), PASTA domain (pYAA049; primers RF584/RF585), or transpeptidase domain (pYAA050; primers RF586/RF587) were deleted by modification of pYAA031 by inverse PCR and subsequent recircularization by ligation.

pYAA051: pYAA048 was further modified to delete the coding sequence of the PASTA domain by inverse PCR with primers RF584/RF585.

### Sporulation efficiency analysis

Overnight cultures of *C. difficile* R20291 were diluted in BHI broth to an OD_600nm_ of 0.01, incubated for 8 h at 37°C, diluted to an OD_600nm_ of 0.0001 and finally incubated overnight. This allowed us to obtain cultures in stationary phase with no detectable spores (T=0). This culture was then incubated for 5 days with vegetative cells and spores enumerated daily. For total viable counts, 10-fold serial dilutions were spotted onto BHIS agar supplemented with 0.1% sodium taurocholate. For total spore counts, the same process was carried out following a 30 min incubation at 65°C. Colonies were counted after 24 h incubation at 37°C and the assay was completed in biological triplicates. Formation of phase bright spores was also followed by phase-contrast microscopy at each time point. Samples fixed in 3.7% paraformaldehyde were imaged using a Nikon Eclipse Ti microscope and analysed using Fiji (Schindelin et al., 2012).

### Microscopy

Bacterial samples were harvested by centrifugation, washed once with PBS and fixed in 4% paraformaldehyde. For phase contrast microscopy, samples were mounted in 80% glycerol and imaged using a Nikon Ti Eclipse inverted microscope. Samples for transmission electron microscopy were fixed as above before additional fixation in 3% glutaraldehyde, 0.1 M cacodylate buffer. Fixed samples were then treated with 1% OsO_4_, dehydrated in ethanol and embedded in araldite resin. Embedded samples were sectioned at 85 nm on a Leica UC6 ultramicrotome, transferred onto coated copper grids, further stained with uranyl acetate and lead citrate and visualized using a FEI Tecnai BioTWIN TEM at 80 kV fitted with a Gatan MS600CW camera.

For fluorescence confocal microscopy, bacteria were grown in TY broth containing 500 nM HADA (Kuru et al., 2015), labelled with 250 nM SNAP-Cell TMR-Star (New England Biolabs) and grown under anaerobic conditions for a further 60 min. Following labelling, cells were harvested at 8,000 x g for 2 min at 4°C and washed twice in 1 ml ice cold PBS. Cells were resuspended in PBS and fixed in a 4% paraformaldehyde at room temperature for 30 min with agitation. Cells were washed three times in 1 ml ice cold PBS, immobilized by drying to a coverslip and mounted in SlowFade Diamond (Thermo Fisher Scientific). Images were captured using a Zeiss AiryScan confocal microscope.

### Antibiotic minimum inhibitory concentrations (MICs)

MICs were determined using the agar dilution method based on the Clinical and Laboratory Standards Institute guidelines (Clinical and Laboratory Standards Institute, 2012). Briefly, agar plates were prepared with a range of concentrations of each antibiotic (2 μg/ml to 1024 μg/ml) using BHI supplemented with defibrinated horse blood (Oxoid) or Brazier’s CCEY agar plates (Bioconnections) supplemented with egg yolk emulsion (Oxoid) and defibrinated horse blood (Oxoid). The plates were dried at room temperature for 2 h and then pre-reduced in the anaerobic cabinet for 2 h. 100 μl of each *C. difficile* strain at an OD_600nm_ of 0.5 was spread onto the plates. The plates were incubated for 48 h in the anaerobic cabinet at 37°C) and MICs were determined by reading the lowest antibiotic concentration on which the bacteria did not grow.

## Results

### *C. difficile* produces a SpoVD homologue that is required for sporulation

The *C. difficile* R20291 genome encodes 10 putative penicillin binding proteins (PBPs) (Table 2) and one predicted monofunctional transglycosylase (CDR20291_2283). In our previous transposon mutagenesis study only two of these, CDR20291_0712 and 0985, were identified as essential for growth *in vitro* (Dembek et al., 2015). However, five of the PBPs were required for formation of heat-resistant spores, including two with homology to the *B. subtilis* cortex specific PBP SpoVD, CDR20291_1067 and 2544. Of these only CDR20291_2544 has the C terminal PASTA domain that is characteristic of the *B. subtilis* sporulation-specific PBPs (Bukowska-Faniband and Hederstedt, 2015). CDR20291_2544 (SpoVD_Cd_) shares 40.1% amino acid identity with *B. subtilis* SpoVD and has the same predicted overall organisation, with an N terminal predicted transmembrane helix, followed by a PBP dimerization domain (PF03717), a transpeptidase domain (PF00905) and the C. terminal PASTA domain (PF03793). *spoVD* is located immediately downstream of CDR20291_2545 (Fig. 2A), encoding a protein with weak homology to *B. subtilis* FtsL (18.8% amino acid identity). Despite the weak similarity, the *C. difficile* and *B. subtilis* proteins are very similar in size (115 and 117 amino acids respectively), have a similar PI (9.57 and 9.63 respectively) and both have a high proportion of lysine residues (22.6% and 14.5% respectively). CDR20291_2545 and *spoVD* appear to be in an operon, with the promoter upstream of CDR20291_2545. In our earlier TraDIS screen, CDR20291_2545 was also found to be required for sporulation, although this may have been due to polar effects on *spoVD*.

**Table 2:** Putative *C. difficile* PBPs.

| *C. difficile* R20291<br>gene designation | Best *B. subtilis*<br>strain 168 hit | Amino acid<br>identity | Essential *in vitro*? |
| --- | --- | --- | --- |
| 0712 | PonA | 27.3% | Yes |
| 2544 | SpoVD | 40.1% | No but required for sporulation |
| 1067 | SpoVD | 27.9%<br>(PbpB 26.6%) | No but required for sporulation |
| 1131 | DacF | 43.8% | No but required for sporulation |
| 1318 | PbpX | 21.3%<br>(PbpE 20.4%) | No |
| 2048 | DacF | 31.5% | No but required for sporulation |
| 0441 | DacF | 30.3% | No |
| 0985 | PbpA | 21.2% | Yes |
| 3056 | PbpX | 20.1% | No but required for sporulation |
| 2390 | DacB | 27.5% | No |

To confirm a role in sporulation, we constructed a clean *spoVD* deletion by homologous recombination and then complemented this mutant by integrating the CDR20291_2545-*spoVD* cassette under the control of the native promoter into the chromosome between the *pyrE* and R20291_0189 genes (referred to here as R20291Δ*spoVD pyrE*∷*spoVD*; Fig. 1A and B). We then analysed the ability of each strain to form heat-resistant spores. In our assay, a stationary phase culture of wild type R20291 gradually accumulated spores, accounting for 81% of the viable counts after 3 days (Fig 1C). In the same assay R20291Δ*spoVD* formed no detectable spores, even after 5 days of incubation (Fig 1D). Complementation completely restored sporulation to wild type levels (Fig 1E). Examination by phase contrast microscopy confirmed the presence of abundant mature phase bright spores in 5 day old cultures of wild type R20291 and the complemented strain R20291Δ*spoVD pyrE*∷*spoVD* (Fig. 2A). In contrast no phase bright objects were observed in cultures of R20291Δ*spoVD*. When visualised at higher magnification using TEM of thin sections, no morphologically normal spores were observed in cultures of R20291Δ*spoVD* (Fig. 2B). Membrane-bound prespores were present, but these structures were irregular in shape and crucially lacked the cortex and protein coat layers seen in R20291 and the complemented strain developing spores. SpoVD is predicted to consist of 3 domains: a PBP dimerization domain, a transpeptidase domain and a PASTA domain (Fig. 3A). To identify which of these were required for viable spore formation, *clip-spoVD* was placed under the control of a constitutive promoter (P_*cwp2*_) in a *C. difficile* expression vector and a panel of mutants, lacking one or more of these domains, were constructed (Table 1). These plasmids were all transferred into R20291Δ*spoVD* and the ability of the expressed SpoVD variant to restore sporulation was evaluated. Only proteins including both the dimerization and transpeptidase domains (SpoVD(DT) or full-length SpoVD(DTP)) restored normal sporulation (Fig. 3B), the PASTA domain was dispensable as observed previously in *B. subtilis* (Bukowska-Faniband and Hederstedt, 2015). This observation was supported by TEM examination, with morphologically normal spores only observed when the full-length or SpoVD(DT) proteins were expressed (not shown).

**Figure 1:**
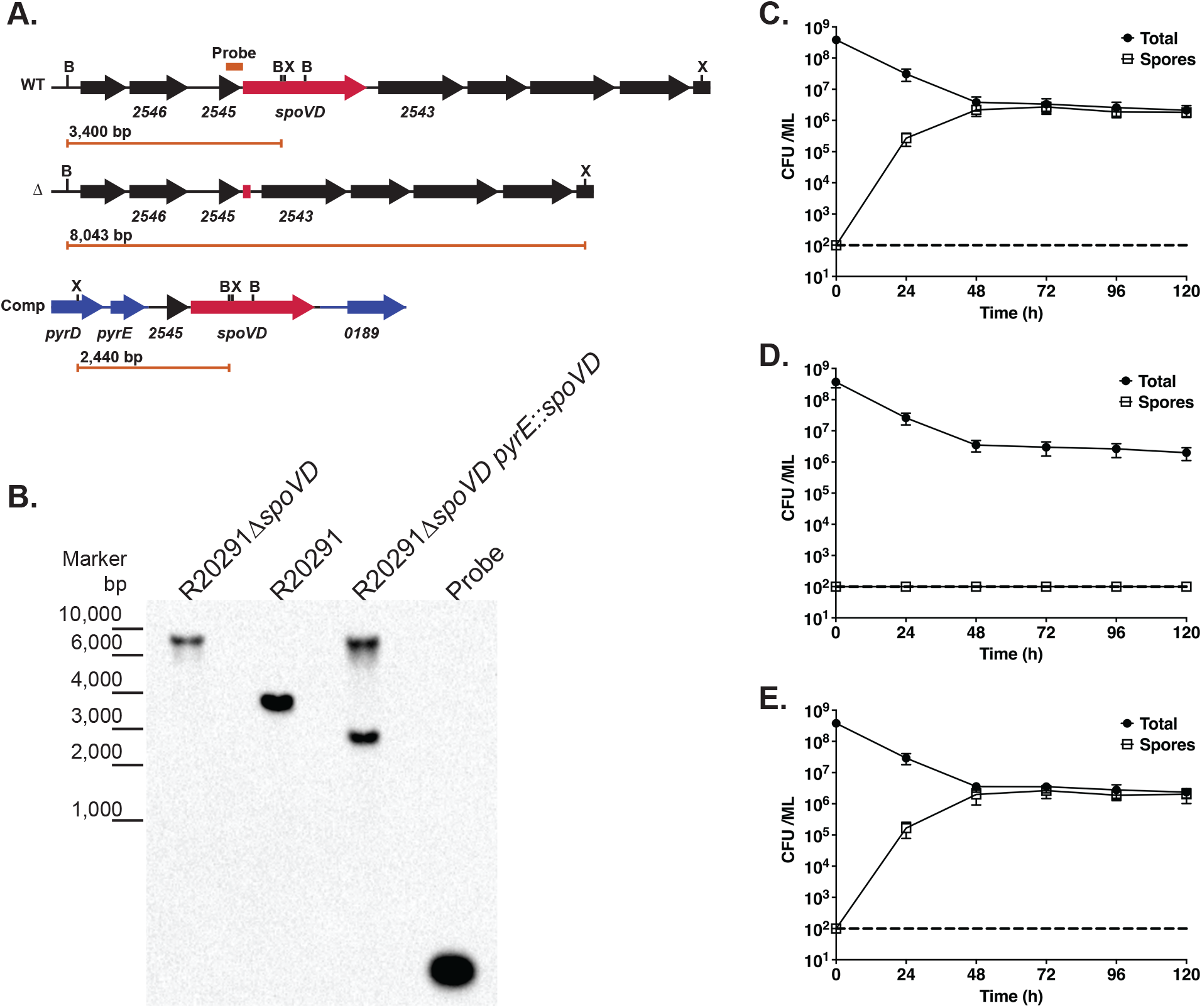
Sporulation requires SpoVD. **A.** Genomic organisation of the native *spoVD* locus (WT), following deletion of the *spoVD* gene (Δ) and following complementation by insertion of R20291_2545 and *spoVD* between the *pyrE* and R20291_0189 genes (Comp). The locations of XmnI (X) and BsrGI (B) sites are indicated, as is the annealing site of the Southern blot probe. The length of the diagnostic restriction product containing the probe sequence is also shown below each locus diagram. **B.** Southern blot analysis of a *spoVD* mutant (R20291Δ*spoVD*), the wild type parental strain (R20291) and complemented strain (R20291Δ*spoVD pyrE*∷*spoVD*). A DNA ladder is shown on the left hand side. The predicted fragment sizes and annealing site of the probe are shown in panel A. **C.-E.** Sporulation efficiencies of the wild type (**C.**), *spoVD* mutant (**D.**) and complemented strains (**E.**). Stationary phase cultures were incubated anaerobically for 5 days with samples taken daily to enumerate total colony forming units (CFUs) and spores, following heat treatment to kill vegetative cells. Experiments were performed in duplicate on biological triplicates with mean and standard deviation shown. The dotted horizontal line indicates the limit of detection of the experiment.

**Figure 2:**
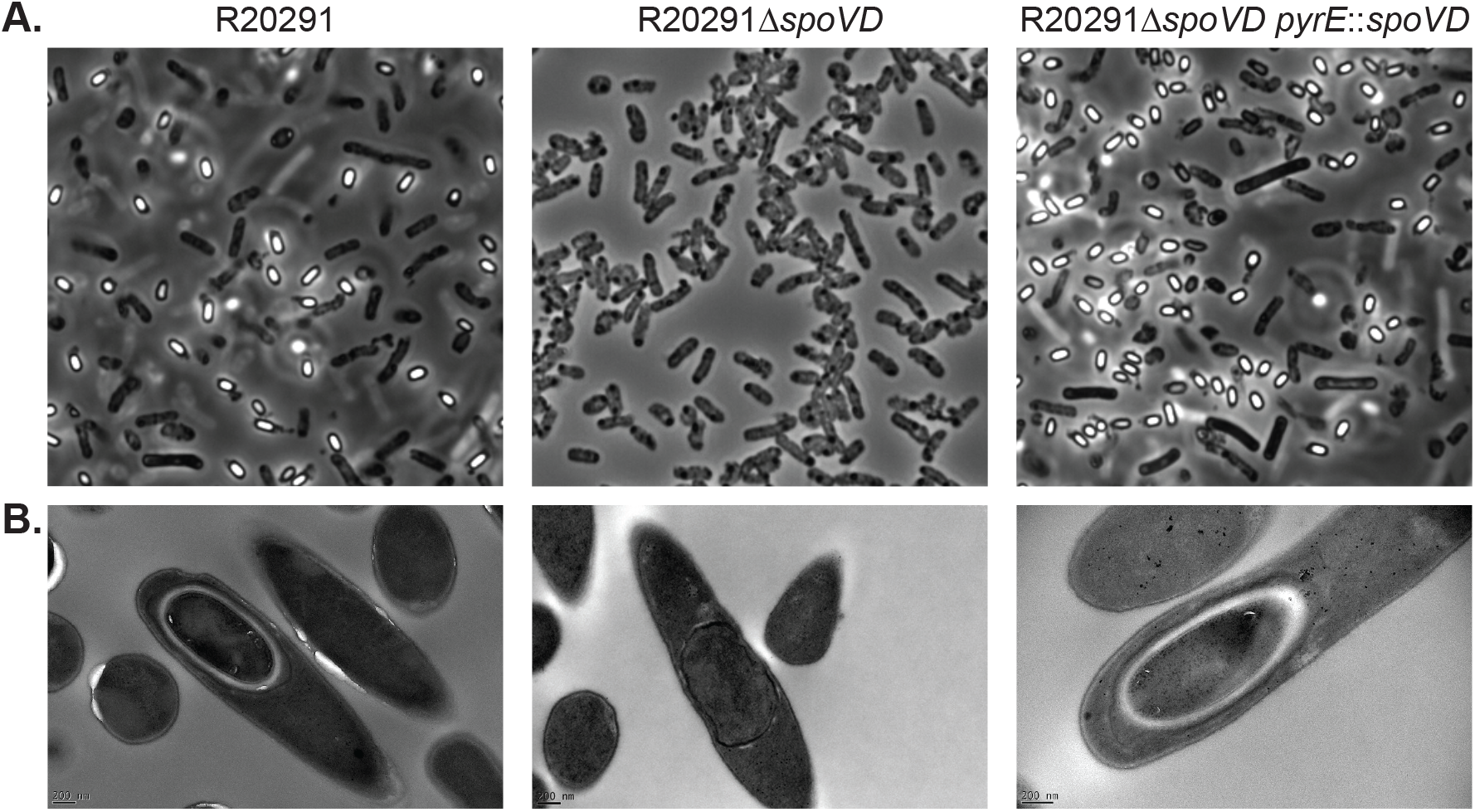
Microscopic analysis of sporulation. Phase contrast light microscopy (**A.**) and negative stained TEM (**B.**) of the wild type parental strain (R20291), *spoVD* mutant (R20291Δ*spoVD*) and complemented strain (R20291Δ*spoVD pyrE*∷*spoVD*). **A.** Cultures were imaged at day 5 of the sporulation assays shown in Figure 1. Spores are visible as ovoid phase bright objects i, while vegetative cells are phase dark bacilli. **B.** TEM imaging of developing spores clearly shows normal spore development in R20291 and R20291Δ*spoVD pyrE*∷*spoVD*; the densely stained core surrounded by a thick, largely unstained cortex layer. Cultures of R20291Δ*spoVD* contained no morphologically normal developing spores, although fully engulfed prespores without a cortex (example shown) were common.

**Figure 3:**
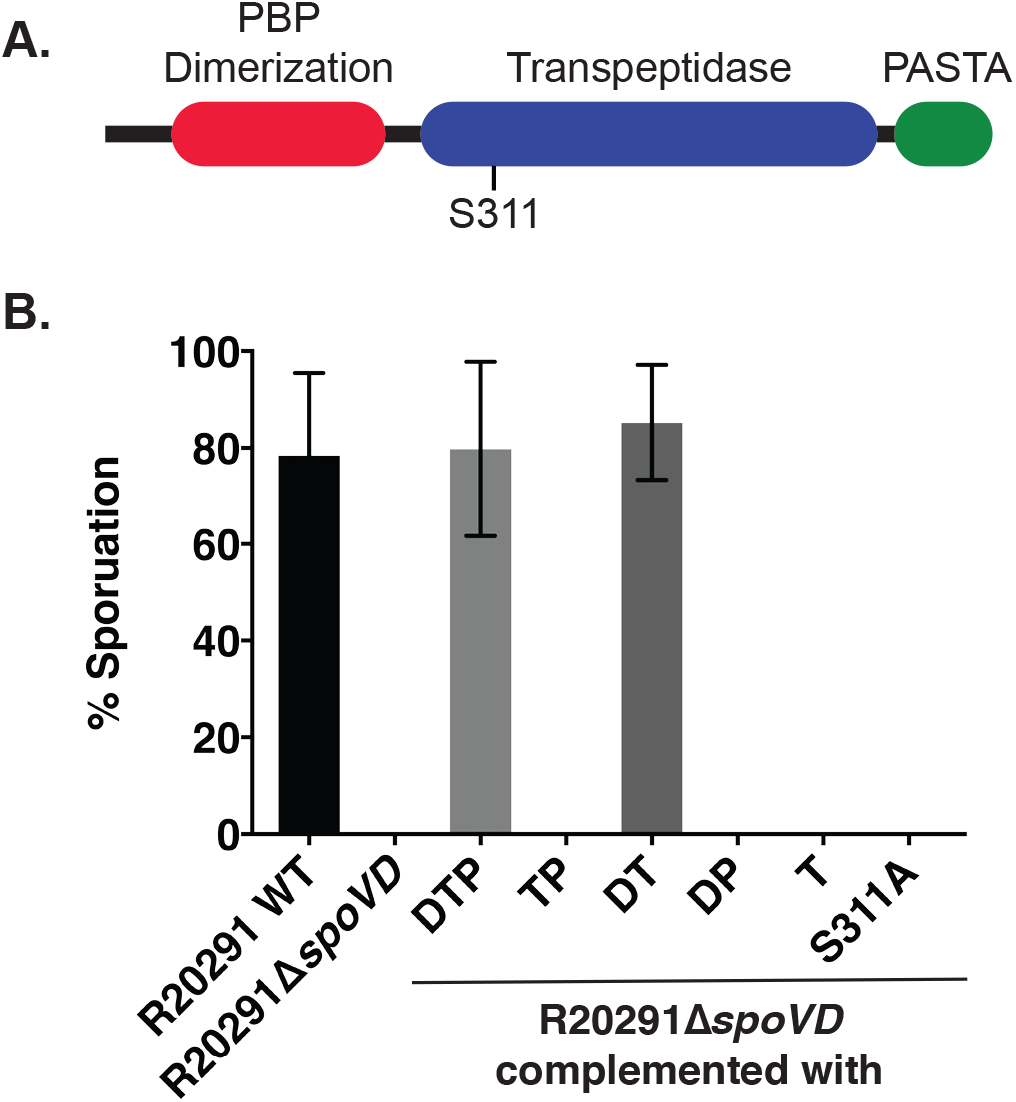
The contribution of SpoVD domains to sporulation. **A.** The domain organisation of SpoVD showing Pfam predictions (El-Gebali et al., 2019). **B.** Sporulation efficiency of R20291, R20291Δ*spoVD* and R20291Δ*spoVD* complemented *in trans* using a plasmid expressing a series of mutant SpoVDs under the control of a constitutive promoter: full-length SpoVD (DTP); SpoVD lacking the PBP dimerization domain (TP), PASTA domain (DT), transpeptidase domain (DP) or both PBP dimerization and PASTA domains (T); SpoVD lacking the active site nucleophile serine (S311A). Shown is the sporulation efficiency after 5 days in broth culture, expressed as number of spores as a percentage of total viable CFUs. Experiments were conducted in duplicate on biological triplicates and mean and standard deviations are shown.

*B. subtilis* SpoVD, and the wider family of class B PBPs, share a conserved active site consisting of 3 non-contiguous motifs that are brought into close proximity in the folded enzyme, SxxK, SxN and KTG (Sauvage et al., 2008). The first of these motifs contains the essential serine nucleophile. SpoVD_Cd_ has all three motifs, with Ser311 as the predicted nucleophile. SpoVD S311A supplied *in trans* was also incapable of complementing the sporulation defect observed in a *spoVD* deletion mutant (Fig. 3B), confirming a role for this residue in cortex synthesis.

### Subcellular localisation of SpoVD

To visualise the cellular localisation of SpoVD, we fused the coding sequence for SNAP to the 5′ end of the *spoVD* gene and transferred this to the *C. difficile* genome in the native locus and under the control of the native promoter. SNAP was then labelled with the fluorescent reagent TMR-Start, while newly synthesised peptidoglycan was labelled with the fluorescent D-amino acid HADA (Kuru et al., 2015). Using Airyscan confocal microscopy we observed weak punctate fluorescence around the periphery of the cell, localizing to the asymmetric division septum once the cell had committed to sporulation (Fig. 4A). Fluorescence then tracked the asymmetric membrane through engulfment (Fig. 4B and C), eventually surrounding the prespore (Fig. 4D). Localization of SNAP-SpoVD clearly preceded significant cortex synthesis as we visualised localisation around the spore without concomitant HADA accumulation (Fig. 4D). Following further spore maturation (Fig. 4E), accumulation of new HADA-labelled peptidoglycan co-localized with SNAP-SpoVD.

**Figure 4:**
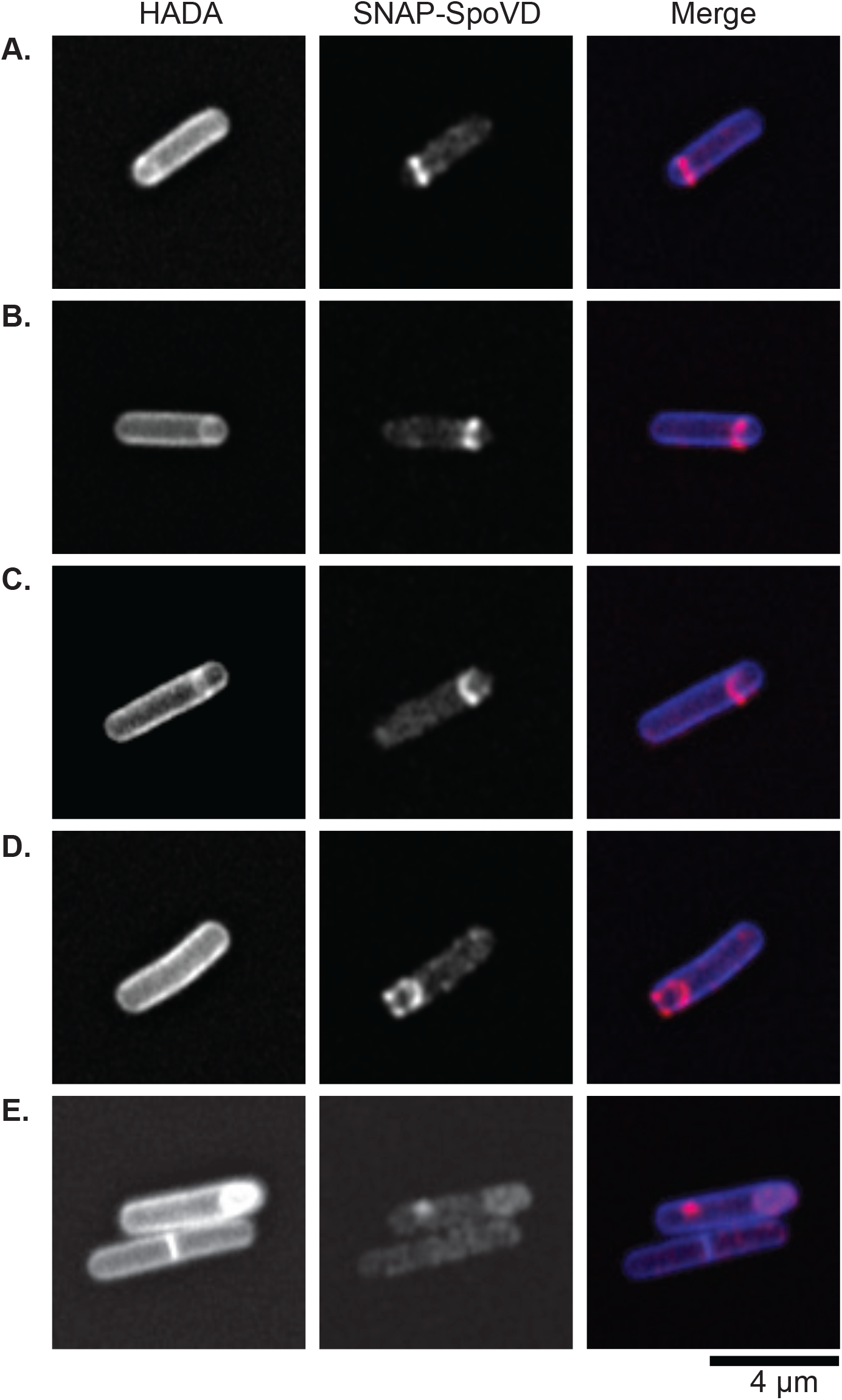
Subcellular localisation of SpoVD. R20291 *snap-spoVD* was grown for 24 h in TY broth containing the fluorescent D-amino acid HADA (500 nM) to label *de novo* synthesised peptidoglycan. The bacteria were then further stained with SNAP-Cell TMR-Star (250 nM) to label SNAP-SpoVD, fixed, mounted in SlowFade Diamond mountant and imaged using a Zeiss AiryScan confocal microscope. Shown are representative cells demonstrating the sequential stages of sporulation: **A.** asymmetric septum placement, **B.**, **C.** and **D.** early, mid and complete prespore engulfment respectively, **E**. spore maturation.

### SpoVD is required for full resistance to cephalosporins

A comparison of the peptidoglycan composition of wild type and Δ*spoVD* cells identified no obvious differences (not shown), suggesting that SpoVD is an exclusive cortex PBP. However, given the extensive complement of PBPs produced by *C. difficile* it is possible that a role in vegetative cell peptidoglycan synthesis could be masked. To determine if SpoVD did contribute to peptidoglycan synthesis more broadly we examined the susceptibility to two PBP-targeting antibiotics, the second- and third-generation cephalosporins cefoxitin and ceftazidime, and the functionally unrelated antibiotic ciprofloxacin. Unsurprisingly deletion of *spoVD* had no effect on ciprofloxacin resistance (Table 3). This mutation did however increase susceptibility to cefoxitin and ceftazidime, decreasing the MIC 4-fold for each, with full resistance restore upon complementation (R20291Δ*spoVD pyrE*∷*spoVD)*. To identify which SpoVD domains were required for this resistance phenotype we complemented the Δ*spoVD* mutant with the same panel of SpoVD truncations described above (Table 3). Interestingly, mutant SpoVDs that lacked either the PBP Dimerization (SpoVD(TP)) or the PASTA domains (SpoVD(DT)) were still fully competent for restoring normal cephalosporin resistance, while that lacking the Transpeptidase domain (SpoVD(DP)) displayed the same aberrant sensitivity as the Δ*spoVD* mutant. This strongly suggests that only the Transpeptidase domain is required for full cephalosporin resistance. To test if the Transpeptidase alone was sufficient for resistance, we then constructed a truncation consisting of this domain alone. However, this construct did not restore resistance to wild type levels. A full-length SpoVD with a single amino acid substitution, replacing the putative nucleophilic serine with alanine (SpoVD(S311A)), also failed to restore resistance, demonstrating that enzymatic activity is required for the observed cephalosporin resistance.

**Table 3:** Antibiotic MICs (μg/ml).

| Strain | Cefoxitin | Ceftazidime | Ciprofloxacin |
| --- | --- | --- | --- |
| R20291 | 128 | 256 | 512 |
| R20291Δ*spoVD* | 32 | 64 | 512 |
| R20291Δ*spoVD pyrE::spoVD* | 128 | 256 | 512 |
| R20291Δ*spoVD*+SpoVD(DTP) | 128 | 256 | n.d. |
| R20291Δ*spoVD*+SpoVD(TP) | 128 | 256 | n.d. |
| R20291Δ*spoVD*+SpoVD(DT) | 128 | 256 | n.d. |
| R20291Δ*spoVD*+SpoVD(DP) | 32 | 64 | n.d. |
| R20291Δ*spoVD*+SpoVD(T) | 32 | 64 | n.d. |
| R20291Δ*spoVD*+SpoVD(S311A) | 32 | 64 | n.d. |

## Discussion

*C. difficile* is the most common cause of hospital acquired infection in the USA and Europe (Magill et al., 2014; Aguado et al., 2015). The formation of a robust spore form is crucial for transmission of infection between patients and for persistence and relapse following treatment (Deakin et al., 2012). However, despite their importance in *C. difficile* pathogenesis, we still know surprisingly little about the underlying molecular mechanisms of sporulation and germination, in part due to a lack of effective genetic tools until recently (Shen, 2019). Much can be learned from the parallels with the well-studied but distantly related species *B. subtilis*, however there are significant differences in the sporulation pathways between the *Bacillales* and *Clostridiales* and even homologous proteins can play subtly different roles (Paredes et al., 2005; Underwood et al., 2009; Fimlaid et al., 2013). Here we have identified and characterised a *C. difficile* homologue of the *B. subtilis* spore cortex PBP SpoVD. We have confirmed that this protein is required for sporulation in *C. difficile* and also identified a surprising role in resistance to cephalosporin antibiotics.

Bioinformatic analysis of the *C. difficile* genome identified 10 genes encoding proteins with significant homology to characterised *B. subtilis* PBPs. In a previous transposon mutagenesis screen we determined that only two of these are essential for growth *in vitro*, but five were required for formation of heat-resistant spores (Dembek et al., 2015). One of these (R20291_2544) encodes a protein sharing 27.9% amino acid identity with *B. subtilis* SpoVD. Despite this relatively weak homology, the two proteins share a similar overall domain organisation and are encoded in a similar genomic context. To determine if this protein played a role in *C. difficile* sporulation we constructed a clean deletion mutant that we found to be incapable of producing viable spores. Microscopic examination of this mutant allowed us to visualise fully engulfed prespores but these structures lacked any obvious cortex. This sporulation defect was fully complemented by integration of *spoVD* (and the upstream R20291_2545 and native promoter) in a distal chromosomal locus. These observations clearly demonstrated that SpoVD plays a crucial role in *C. difficile* sporulation and is required for the synthesis of cortex peptidoglycan. We then demonstrated that the sporulation defect in a *spoVD* mutant could be complemented by expression *in trans* of a mutant SpoVD lacking the C terminal PASTA domain but that mutation of the PBP dimerization or transpeptidase domains resulted in a non-functional SpoVD. This is in full agreement with previous *B. subtilis* studies that showed that the PASTA domain was dispensable for cortex synthesis (Bukowska-Faniband and Hederstedt, 2015). By comparison with the *B. subtilis* sequence we were also able to putatively identify the active site nucleophile serine as S311 and confirmed this role by mutation to alanine, resulting in a non-functional SpoVD.

It has been shown previously that *B subtilis* SpoVD localises to the asymmetric septum upon initiation of sporulation and ultimately to the developing spore following engulfment (Sidarta et al., 2018). To visualise this process in *C. difficile* we generated a strain expressing SNAP-SpoVD under the control of the native promoter. Super-resolution fluorescence microscopy imaging of this strain showed clear localisation of SNAP-SpoVD to the asymmetric septum and to the developing spore. Intriguingly we also observed weak punctate fluorescence staining around the periphery of the mother cell. This could be indicative of mislocalisation as a result of the N terminal SNAP fusion or could suggest a broader role for SpoVD in vegetative cell peptidoglycan synthesis. To test this latter possibility, we examined the peptidoglycan composition of wild type and *spoVD* mutant cells but observed no obvious differences. Given the enormous potential for redundancy with 10 encoded PBPs it is possible that small differences could be missed in this analysis. To examine more subtle effects, we analysed resistance to two PBP-targeting cephalosporin antibiotics, cefoxitin and ceftazidime. Surprisingly the *spoVD* mutant displayed a 4-fold reduction in MIC to both antibiotics. Although small, this difference in MIC was reproducible and complemented perfectly when *spoVD* was added back. This unexpected observation strongly suggests that SpoVD is active in vegetative cells and is not spore-specific as has been suggested for *B. subtilis*.

Sporulation of *C. difficile* represents one of the most pressing clinical challenges in tackling recurrent disease in individual patients as well as preventing outbreaks in nosocomial settings. However, this cell differentiation pathway also represents a promising target for the development of *C. difficile*-specific therapeutics. In order to exploit this potential we must first develop a deeper understanding of both the complex regulatory processes that underpin sporulation as well as the function of the effector proteins that direct differentiation. Here we have identified and characterised a PBP that is absolutely required for production of viable spores and that we believe is a promising target for future therapeutics aimed at preventing recurrent disease and transmission.

## Acknowledgements

We would like to thank Darren Robinson and Christa Walther (The Wolfson Light Microscopy Facility, University of Sheffield) for their light microscopy support and training, Stephane Mesnage (University of Sheffield) for assistance with peptidoglycan analysis, Chris Hill (Electron Microscopy Unit, University of Sheffield) for thin sectioning and transmission electron microscopy, Nigel Minton (University of Nottingham) for supplying plasmids for homologous recombination, Adriano Henriques for supplying pFT46, and Simon Jones and Shuwen Ma (University of Sheffield) for synthesis of HADA.

## Author contributions

Y.A.A. and R.P.F. designed and coordinated the study. Y.A.A., P.O. and J.A.K. performed the experiments. R.P.F. wrote the paper with input from all co-authors.

